# Coordinated alterations in SWI/SNF chromatin remodeling complex, mRNA splicing and mRNA methylation as hallmarks of salivary gland adenoid cystic carcinoma (SACC)

**DOI:** 10.64898/2026.10.07.757230

**Authors:** Katarzyna Stawarz, Natalia Rusetska, Hubert Salamaga, Szymon Kubala, Irena Paczkowska, Pawel Cwiek, Alicja Berezowska, Maryam Jozghorbani, Jakkapong Kluebsoongnoen, Magdalena Wilga, Susanne Poveda, Agnieszka Swic, Monika Durzynska, Adam Galazka, Ewa Chmielik, Tomasz Rutkowski, Andrzej Kawecki, Beata Jagielska, Piotr Zielenkiewicz, Elzbieta Sarnowska, Tomasz J. Sarnowski

## Abstract

Salivary gland adenoid cystic carcinoma (SACC) is a rare, poorly understood malignancy. Here, using spatial transcriptomics, we show that disturbances in epigenetic regulation including chromatin remodeling, epitranscriptomic control by m6A RNA methylation, and RNA splicing may contribute to SACC. We demonstrated that machineries responsible for these processes-SWI/SNF chromatin remodeling complex (CRC), NineTeen Complex (NTC), and m6A RNA methylation complex interact directly. Immunohistochemistry on 48 SACC tumor samples indicated altered abundance of METTL3, METTL14, WTAP (m6A RNA methylation) and PRP19, CDC5L, PLRG1 (NTC-splicing) proteins. Chromatin immunoprecipitation indicated direct regulation of genes encoding subunits of SWI/SNF, NTC and RNA methylation complexes, and some EMT (epithelial-to-mesenhymal) marker genes by BRM (BRAHMA). Transcriptomic re-analysis revealed broad alterations in transcript usage, and splicing events in SACC. Collectively, our findings provide the first evidence that coordinated disruption of chromatin remodeling, RNA methylation, and splicing contributes to SACC development and progression, highlighting new avenues for therapeutic exploration.

## Main

<u>S</u>alivary gland <u>a</u>denoid <u>c</u>ystic <u>c</u>arcinoma (SACC) is a rare neoplasm characterized by relatively slow growth, frequent local recurrences, and a propensity for distant metastases [1]. Beside surgery, current therapeutic modalities including radiotherapy, chemotherapy, or immunotherapy, appear ineffective reflected in estimated 10-year overall survival rates at 37%-65% [2]. Given its rarity in clinic and some previous preclinical studies based on contaminated cell lines [3], SACC remains inadequately understood, underscoring the imperative to elucidate its foundational pathogenesis and molecular changes.

Disruptions in regulatory processes, such as alterations in chromatin structure, RNA methylation or splicing, have been implicated in the development of various cancers [4–6]. The SWI/SNF ATP-dependent chromatin remodeling complexes (CRCs) alter chromatin nucleosomal structure to modulate DNA accessibility, directly influencing transcriptional elongation by RNA polymerase II (RNAPII). The kinetics of RNAPII elongation, co-transcriptional splicing and chromatin structure are interdependent [7] and influenced by the functional SWI/SNF CRCs [8,9]. Specific subunits of the SWI/SNF CRCs, such as BRM/SMARCA2 and BRG1/SMARCA4, have been shown to interact with splicing regulators, thereby directly influencing splicing decisions, with far more profound role of BRG1 in this process [9,10]. Additionally, mutations or disruptions in genes encoding SWI/SNF subunits, such as BRG1/SMARCA4 and BAF250A/ARID1A, are frequently observed in cancers and associated with their own splicing deregulation. The changes in splicing of SWI/SNF subunits encoding genes contribute to tumor progression by altering exon inclusion or skipping, leading to the production of oncogenic isoforms [11]. SWI/SNF-mediated chromatin remodeling influences histone modifications, which are known to affect splicing outcomes. These epigenetic changes alter chromatin environment near splice sites, affecting recruitment of splicing machinery [12,13]. The NineTeen Complex (NTC), is spliceosomal complex required for catalytic spliceosome activation and efficient pre-mRNA splicing. Increasing evidence link aberrations in NTC components with cancer development and progression through effects on RNA splicing, genome maintenance, and pathogenic splice isoforms generation [14,15]. Dysregulation of NTC components impairs homologous recombination repair, leading to genomic instability, a hallmark of cancer [16].

The N6-methyladenosine (m6A) methyltransferase complex, commonly referred to as the m6A “writer” complex, is a key regulator of the dynamic deposition of m6A, the most abundant internal modification of eukaryotic mRNA. This epitranscriptomic mark influences multiple aspects of RNA metabolism, including splicing, stability, translation, and decay. Increasing evidence indicates that dysregulation of m6A deposition contributes to cancer initiation and progression. The core m6A writer complex is composed of METTL3 (catalytic subunit); METTL14, which facilitates substrate recognition; and WTAP (Wilms Tumor 1-Associated Protein), stabilizing the METTL3– METTL14 heterodimer and promoting its recruitment to specific RNA targets. Functionally, aberrant m6A deposition can promote malignant phenotypes by increasing the stability or translational efficiency of oncogenic mRNAs [17] while in other contexts it may accelerate the decay of tumor suppressor transcripts [18]. m6A-dependent regulation of stemness-related genes has been implicated in cancer cells supporting self-renewal and chemoresistance in cancer [19,20].

Emerging evidence suggests that SACC is shaped by extensive epigenetic deregulation, marked by impairment of SWI/SNF CRCs [21,22] and by aberrant splicing programs [23,24].

Here, we define a molecular framework underlying key features of SACC. We show that mRNA splicing, SWI/SNF CRCs and RNA methylation machinery are functionally interconnected and coordinately disrupted in SACC. The detailed analysis using spatial transcriptomics, immunohistochemistry and deep transcriptomic study uncovered molecular differences between SACC with and without nerve invasion. We found that the alterations in the presence of SWI/SNF, NTC and RNA methyltransferase complex are more pronounced when the invasion of SACC on nerves occurs. We deduced that the impairment of SWI/SNF CRC may be a causative factor for alterations observed in SACC. Thus, our study indicate that the impairment of fine-tuning of SWI/SNF-dependent chromatin remodeling, mRNA splicing and mRNA methylation may be one of the most important factors in SACC progression.

## Results

### SWI/SNF, NTC and m6A mRNA methylation complex subunits expression correlates with BRM overexpression in SACC and may depend on cancer stage

We have previously shown that SACC is featured by the altered balance between SWI/SNF CRCs subclasses towards a subclass containing BRM ATPase [21] what is in line with altered transcriptional profile of this cancer and its molecular characteristics (e.g. chemoresistance, affected splicing, chromatin remodeling, enhanced proliferation). These findings did not provide exhaustive explanation for the occurrence of characteristic for SACC pronounced alterations in transcript formation like loss of alternative 3’ acceptor, alternative transcription termination and alternative 5ʹ acceptor sites, enhanced multiple exon skipping or alternative transcription start sites usage [24].

To address this question and to integrate the transcriptomic data with SACC tissue morphology we employed the 10x Genomics Visium Spatial Transcriptomics platform and captured spatially-resolved gene expression in SACC samples (Fig.S1A). The 2 SACC samples were analyzed, first representing early stage of cancer without nerve invasion, second-advanced stage of SACC with strong nerve invasion (Fig. S1B). Both samples strongly differed in gene expression patterns (Fig.S1C).

In less advanced SACC, the pattern of *SMARCA2* transcript encoding BRM ATPase subunit of SWI/SNF CRCs corresponded to BRM protein abundance and strongly correlated with expression of genes encoding other SWI/SNF subunits (*ARID1A*, *SMARCA4*, *SMARCC1*, and *SMARCB1*). Only *SMARCC2* exhibited much broader expression in this sample. The expression of *METTL3*, *METTL14* and *CDC5L* demonstrated the same pattern like BRM. Only *PLRG1* expression was much broader and resembled *SMARCC2* pattern (Fig.1A). In advanced SACC sample (with neural invasion), the *SMARCA2* transcription and BRM abundance patterns were not fully corresponding indicative of differential regulation at transcript and protein levels. The expression pattern of other SWI/SNF complex subunits and *METTL3*, *METTL14* as well as *CDC5L* and *PLRG1* were similar. Of note, in more advanced SACC stage, the *ARID1A*, *SMARCA2* and *PLRG1* expression had nearly identical patterns resembling *SMARCB1*, *SMARCC1*, *METTL3* and *CDC5L* mRNA abundance (Fig.1B). These results suggest the existence of coordinate deregulation of SWI/SNF CRC, NTC and m6A mRNA methyltransferase complexes likely leading to dramatic transcriptomic changes observed in advanced SACC.

**Figure 1.**
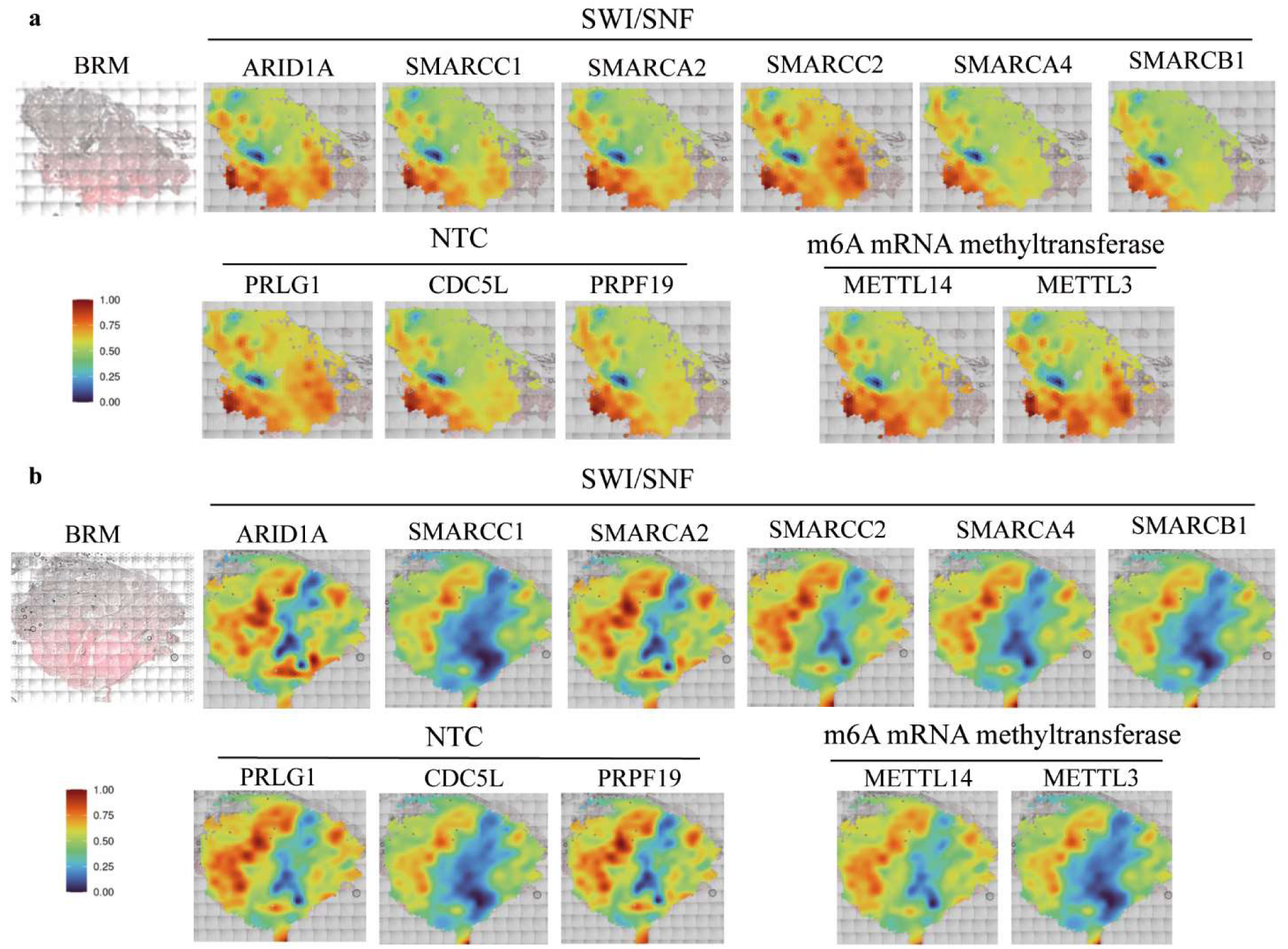
The spatial transcriptomics analysis of patients’ samples with and without invasion on nerves. (a) The BRM abundance in SACC patient sample without neural invasion, and spatial transcriptomics analysis for BRM encoding gene SMARCA2, other subunits of SWI/SNF complex like ARID1A, SMARCA4, SMARCC1, SMARCC2 and SMARCB1 and NTC subunits: PLRG1, CDC5L and m6A mRNA methylation complex: METTL3 and METTL14. (b) The BRM abundance in SACC patient sample with advanced stage –present neural invasion, and spatial transcriptomics analysis for BRM encoding gene SMARCA2, other subunits of SWI/SNF complex like ARID1A, SMARCA4, SMARCC1, SMARCC2 and SMARCB1 and NTC subunits: PLRG1, CDC5L and m6A mRNA methylation complex: METTL3 and METTL14.

### Subunits of the SWI/SNF CRCs interact with NTC splicing and N6-methyladenosine (m6A) methyltransferase complexes

Given our spatial transcriptomics analysis indicated potential interplay between chromatin remodeling, RNA splicing and RNA methylation, we investigated the interaction between BAF155 (a core subunit of the SWI/SNF CRC) and CDC5L (a component of the NineTeen Complex involved in pre-mRNA splicing) using co-immunoprecipitation (IP) and yeast two-hybrid (Y2H) assays. Our results revealed that BAF155 a SWI/SNF subunit interacts with CDC5L a NTC subunit in both yeast-two-hybrid assay (Fig.2A) and in direct immunoprecipitation from HeLa cells nuclei (Fig.2B), indicative of that the interaction of SWI/SNF and NTC complexes is not only limited to BRM and BRG1 ATPases as it has been proposed earlier [10]. As the alternative splicing events frequently depend on the mRNA methylation [25,26] we tested if the m6A methyltransferase complex may interact with the SWI/SNF-NTC system and confirmed that both NTC and SWI/SNF complexes’ subunits interact with WTAP and METTL3 proteins, the components of m6A mRNA methyltransferase complex (Fig.2C). Collectively, our data clearly indicate that chromatin remodeling, transcription, RNA splicing and mRNA modifications may occur spatio-temporary. Such interaction may serve as attractive basis for explanation of transcriptional changes characteristic for SACC.

**Figure 2.**
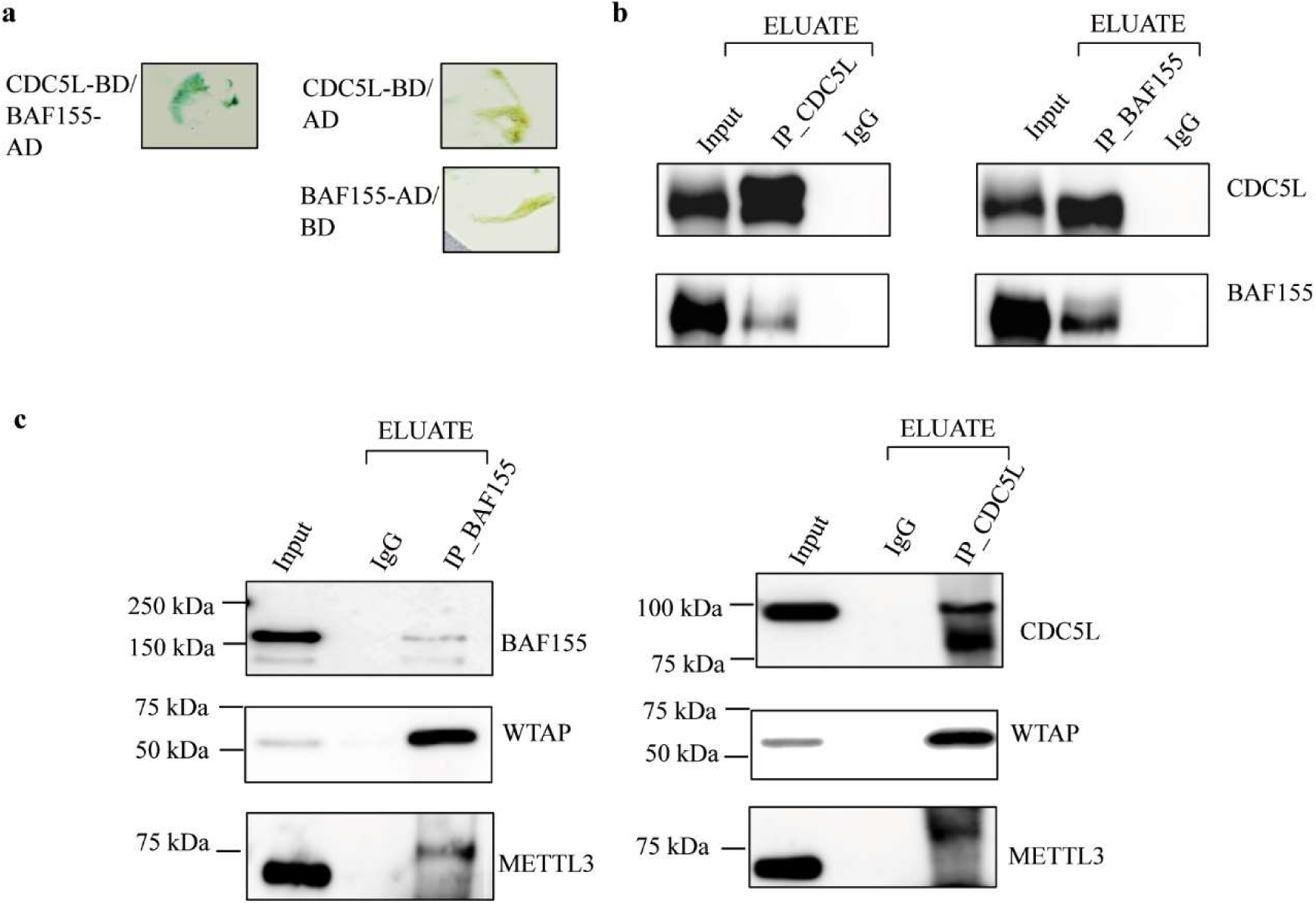
SWI/SNF chromatin remodeling complex, NTC and m6A mRNA methylation machinery components interact in the cell nuclei. (a) yeast-two-hybrid assay indicates interaction between BAF155 SWI/SNF subunit and CDC5L a core NTC component; (b) co-immunoprecipitation from HeLa cell nuclei using anti-BAF155 and anti-CDC5L antibodies confirms that SWI/SNF-NTC interactions are not limited to the ATPase subunits; (c) co-IP using anti-BAF155 and anti-CDC5L together with m6A mRNA methylation complex subunits WTAP and METTL3 indicate interactions between SWI/SNF, NTC and m6A mRNA methylation complexes.

### BRM abundance influences expression of NTC and m6A RNA methylation complex subunits and EMT markers in SACC

To verify the interdependence of BRM and expression of NineTeen and m6A mRNA methyltransferase complexes’ subunits we developed the tetracycline inducible SACC cell line UM-HACC-2A with downregulation of BRM ATPase encoding *SMARCA2* gene using shRNA, specifically targeting ATPase domain-encoding region of BRM protein. We selected two independent clones for analysis. In both clones No.1 and No.2 the downregulation of BRM was observed (Fig.3A). Similarly, the discrete downregulation of protein levels of CDC5L a NTC subunit and WTAP and METTL3 - a m6A RNA methyltransferase complex subunits, was found in the BRM-depleted cell line (Fig.3A). Consistently, the qRT-PCR indicated the downregulation of SMARCA2/BRM gene transcript although, for clone No.2, the less robust downregulation of SMARCA2/BRM transcript level was observed comparing to clone No.1. In clone No.1 the stronger effect on transcript level of *METTL3* was observed (Fig.2B) for other examined genes: WTAP, CDC5L and PLRG1 the transcript level was downregulated similarly for both clones (Fig.3B). We subsequently tested if the depletion of BRM has any effect on the aggressiveness of SACC cells. We therefore performed the Western blot assessment for N-cadherin and vimentin and found that BRM depletion was associated with decreased protein abundance of both EMT markers (Fig. 3C).

**Figure 3.**
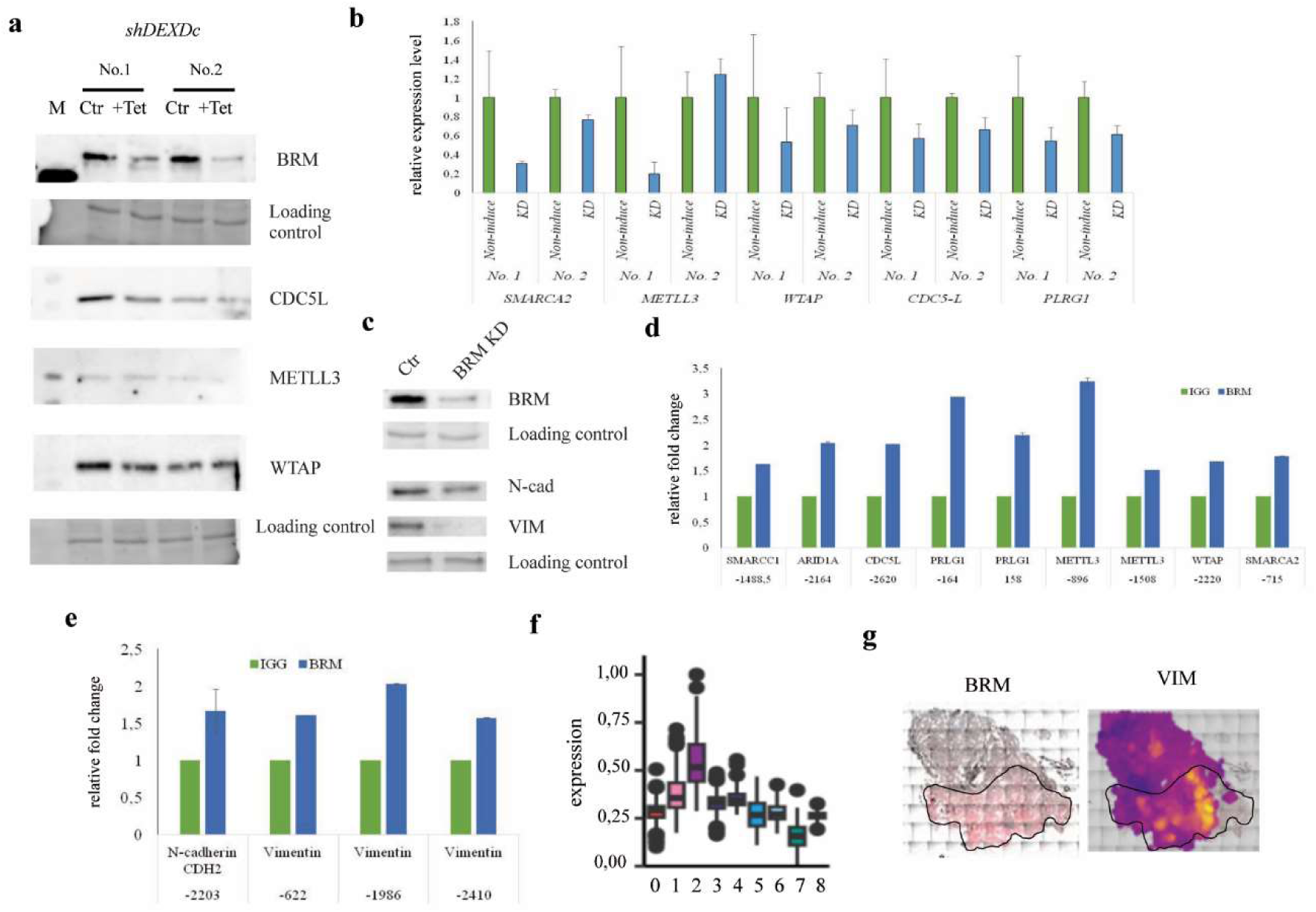
BRM directly controls expression of SWI/SNF, NTC and m6A methyltransferase complexes’ subunits and EMT markers in SACC. The two independent SACC cell lines carrying tetracycline-inducible shRNA (shDEXDc) targeting *SMARCA2* gene exhibit downregulation of NTC and m6A methyltransferasease complexes. (a) The Western blot using antibody against BRM, WTAP, CDC5L, and METTL3 indicated downregulation of BRM in tetracycline inducible lines compared to non-induced control, as well as discrete downregulation of all examined proteins. 0,5% TCE staining was used for visualization of random band serving as a loading control. (b) The qRT-PCR using targeted genes specific primers on tetracycline inducible cell lines was performed, and indicated the downregulation of *SMARCA2* and other examined genes, except METTL3 in clone No.2 The three independent biological replicates were performed on each line and the average value was indicated on graph. (c) The Western blot using antibody against N-cadherin and vimentin indicated decreased abundance of these EMT markers when BRM is downregulated. 0,5% TCA staining was used for visualization of random band serving as a loading control. (d) The chromatin immunoprecipitation indicated direct binding of BRM to its target *loci* encoding NTC and m6A methyltransferase complexes. (e) BRM directly binds the EMT markers encoding genes. (f) The vimentin expression in different clusters in SACC patient sample. (g) The spatial distribution of VIM transcript and BRM protein abundance.

### BRM binds *loci* of genes encoding subunits of NTC, m6A methyltransferase complexes and epithelial to mesenchymal transition markers

Given the de-regulation of genes encoding NTC and m6A RNA methyltransferase complexes as well as EMT markers in the BRM-downregulated SACC cell line we tested if these *loci* may be directly targeted by the BRM ATPase of SWI/SNF CRCs. We found BRM occupancy on such genes encoding subunits of NTC complex as CDC5L (-2666 to 2574bp from TSS, fold change 2.02) and PLRG1 (+101 to +216bp and -221 to -107 from TSS, fold change 2.19 and 2.96 respectively) and m6A methyltransferase complex-METTL3 (-947 to -846bp and -1553 to -1463bp from TSS, fold change 3.24 and 1.52 respectively), and WTAP (-2277 to – 2163 from TSS, fold change 1.68). Interestingly, we found BRM occupancy on the promoter regions of following genes encoding subunits of SWI/SNF CRCsː SMARCA2 (-765 to -665bp from TSS, fold change 1.78), SMARCB1 (-2865 to -2761bp from TSS, fold change 1.55), SMARCC1 (-1540 to -1437bp from TSS, fold change 1.62) and ARID1A (-2209 to -2119bp from TSS, fold change 2.03). Of note, BRM targeted additionally promoter regions of genes encoding EMT markersː Vimentin (-667 to -578bp; -2033 to -1939bp; -2451 to -2369bp from TSS, fold change 1.61, 2.03 and 1.57 respectively) and N-cadherin (-2240 to -2158bp from TSS, fold change 1.67) (Fig. 3D, E).

Given this observation we further explored the spatial transcriptomics datasets and found significant enrichment in the epithelial-to-mesenhymal transition-related genes in both patients, however it was more pronounced in patient without invasion on nerves, e.g. present in more clusters, main EMT gene, Vimentin, was upregulated in regions where BRM protein occurred (Fig. 3F, G Fig. S2 A, B). Given potential BRM regulatory role in the maintenance of functionality of SWI/SNF/NTC/m6A RNA methyltransferase module and in the aggressiveness of SACC by targeting mainly distal promoter regions we performed mapping of transcription factors and other DNA binding/modifying proteins which may target regions bound by BRM in these *loci* with special focus on BRM or SWI/SNF complex interactors (Supplementary table 1) and we found factors involved in transcription activation, repression, m6A machinery recruitment and control of alternative transcription.

### Subunits of the NTC complex are overexpressed and mRNA methylation complex components are deregulated in SACC samples

Our previous study revealed deregulation of processes related to the formation and regulation of RNA, including mRNA splicing, in SACC cells and indicated elevated abundance of BRM ATPase of SWI/SNF complexes in this cancer [21]. This observation together with direct BRM binding of *loci* encoding NTC and m6A methyltransferase complexes prompted us to check if the expression of NTC complex subunit is altered in SACC samples using IHC staining and demonstrated an increased amount of PRP19 and PLRG1 proteins in SACC cells, with only a discrete increase in the amount of the CDC5L subunit (Fig.4A). In 5 cases the membrane staining for CDC5L was observed (Fig.4B).

**Figure 4.**
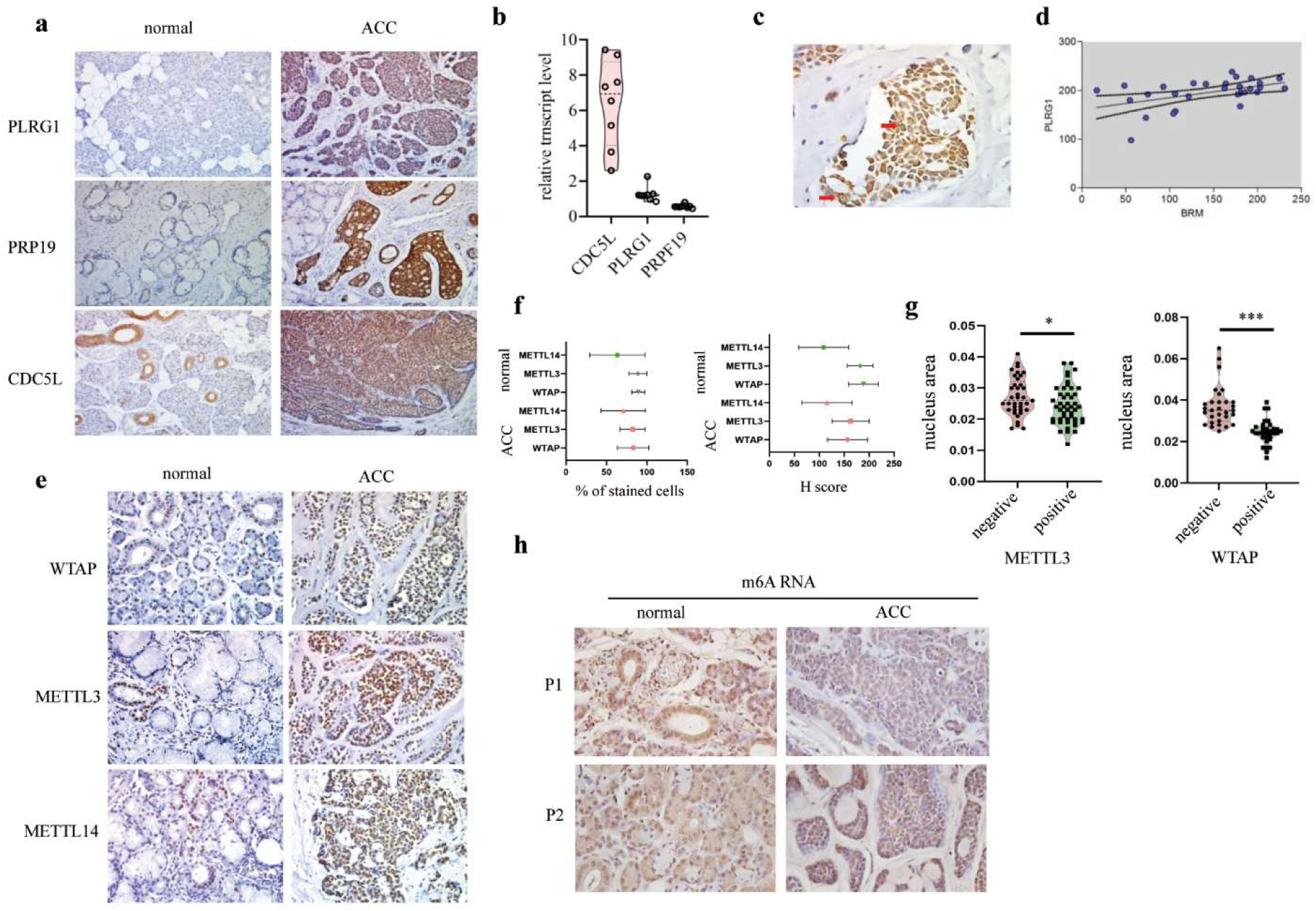
SACC exhibits deregulation of NTC complex subunits and altered abundance of m6A RNA metyltransferase complex subunits. (a) IHC analysis of the presence of the NTC splicing complex subunits-PLRG1, PRP19 and CDC5L (magnification x200). (b) Analysis of the mRNA level for the genes encoding the subunits of the NTC complex PRP19, CDC5L and PLRG1. (c) Membrane staining for CDC5L in SACC cells (magnification x630). * p-value<0.05. (d) Positive correlation between the amount of BRM ATPase and the PLRG1 protein. Correlation acc. Pearson, Pearson r = 0.4858, p-value = 0.0075. (e) IHC analysis of the subunits of the m6A mRNA methylation complex -WTAP, METTL3 and METTL14 (magnification x200). (f) % of stained cells and H-score analysis of all examined proteins in healthy and tumor tissue. (g) Analysis of the METTL3 and WTAP positive and negative nuclei in cancer cells. (h) Analysis of m6A mRNA in normal and SACC cells in salivary gland indicates decreased m6A levels in cancer (magnification x400); * p-value<0.05, *** p-value<0,0005.

Transcriptomic analysis revealed reduced expression of genes encoding PRP19 and PLRG1 in SACC cells, while CDC5L gene expression was increased in tumor cells (Fig. 4C), indicative of multilevel deregulation of NTC in SACC. The amount of PLRG1 protein in SACC positively correlated with BRM protein levels (Fig.4D).

Given our observation that BRM directly binds the genes encoding m6A mRNA methyltransferase complex and their alternative splicing is altered in SACC samples we tested their protein levels in SACC. The analysis revealed no significant difference in the percentage of cells stained for WTAP, METTL3, and METTL14 proteins between tumor and healthy tissues (Fig. 4E, Fig. 4F). However, comparative evaluation of protein levels across all tested markers demonstrated a significant reduction in H-score values for WTAP and METTL3 in tumor samples (p < 0.05), suggesting decreased staining intensity in cancer tissue compared to healthy control.

Further examination of SACC cell nuclei positive or negative for WTAP and METTL3 staining revealed that nuclei expressing detectable levels of these m6A complex subunits were significantly smaller than those without staining (Fig. 4G). Additionally, m6A mRNA analysis using an anti-m6A antibody showed a markedly lower abundance of methylated mRNA in SACC tissues compared to healthy controls (Fig. 4H), reinforcing the observation of dysregulated m6A modification in SACC samples.

### Alternative splicing aberrations in SACC samples

The intriguing discrepancy between protein and transcript levels for some genes (e.g. encoding subunits of NTC splicing machinery) in SACC prompted us to investigate the alterations in splicing in this cancer. We thus analyzed splicing variants using RNA-seq datasets from publicly available databases and identified 8996 isoforms undergoing at least one isoform switch accounting to 15 122 isoform switch events affecting 5498 genes in SACC samples comparing to healthy tissue (Fig. 5A). These splicing alterations resulted in significant consequences, including complete ORF loss, domain gain, and intron retention (Fig. 5B). Additionally, we observed various splicing aberrations, such as alternative 5’ splice site (A5) gain or loss, intron retention (IR) gain, and alternative transcription start site (ATSS) gain or loss (Fig. 5C). Notably, the most frequent splicing abnormality in SACC was alternative transcription termination site (ATTS) gain or loss (Fig. 5C). Furthermore, a shift in splicing isoform usage was detected, particularly in ATSS and ATTS isoforms, which were predominantly utilized in SACC cells (Fig. 5D). Importantly, cancer cells exhibited distinct isoform usage patterns than healthy cells (Fig. 5E).

**Figure 5.**
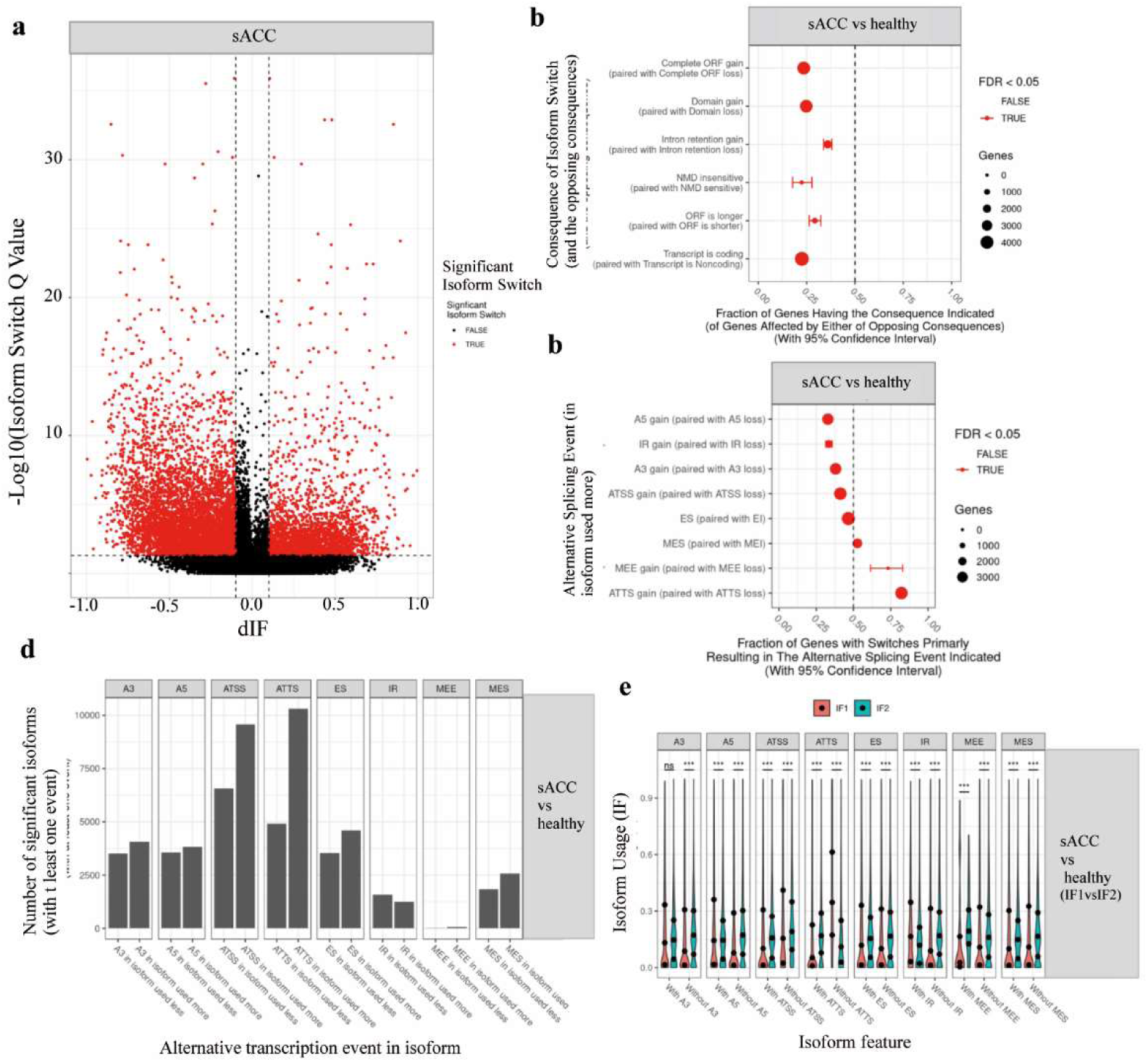
SACC is characterized by global changes in RNA splicing. (a) The analysis of alternatively spliced genes in SACC samples comparing to normal healthy salivary gland tissue indicates increased number of isoforms switch in SACC. (b) Fraction of genes alternatively spliced which having a consequences. (c) Fractions of genes with switched splicing isoforms. (d) Numbers of significant alternatively spliced isoforms. (e) The distribution of used spliced isoforms in SACC comparing to healthy tissue.

Among genes with affected alternative splicing in SACC we found genes encoding main subunits of SWI/SNF CRCs, namely *SMARCA2*, *SMARCA4*, *SMARCB1* and *SMARCC1* (Supplementary Fig. 3-6) further supporting deregulation of this machinery in SACC.

We found that the SACC exhibits characteristic alterations in the transcript profile for PRP19, and PLRG1 subunits of NTC demonstrated by the transcript balance movement towards protein coding classes of transcripts (Fig. S7 and Fig. S8) collectively indicating possible enhanced/aberrant NTC activity in SACC.

Gene encoding METTL3 subunit of m6A RNA methyltransferase complex exhibited distinct splicing isoforms in SACC comparing to healthy salivary gland tissue (Fig. S9A). While the total expression level of the METTL3 gene remained unchanged between SACC samples and healthy controls, a significant overexpression of both the main METTL3 isoform and an NMD-sensitive (Nonsense-Mediated mRNA Decay) isoform was observed (Fig. S9B). This dysregulation may further support observed by us alteration in m6A RNA methyltransferase complex subunits in SACC patient samples and indicate impairment of m6A writer machinery in this cancer type (Fig. 4 E-H).

## Discussion

Molecular alterations in complexes regulating mRNA methylation, chromatin nucleosomal structure, and mRNA splicing are often linked to cancer development and progression [27,28].

The expression of genes encoding WTAP, METTL3, METTL14 m6A mRNA methyltransferase, various subunits of SWI/SNF and NTC complexes, as well as their corresponding protein levels in SACC remained largely unexplored. Here, we demonstrated that SACC exhibits aberrant abundance of these machineries. Numerous studies have demonstrated that dynamic changes in RNA methylation at the adenosine residue are intricately linked to cancer development [29–31]. It has been established that, in the absence of WTAP subunit of the mRNA methyltransferase complex, the level of m6A decreases [32]. In various cancer types, another subunit, METTL3, has been implicated in promoting cancer progression, functioning as an oncogene [33]. However, in certain malignancies, it assumes a suppressor role [34] but in other cancer types reduced METTL3 activate the AKT pathway, thereby enhancing cell proliferation and aggressiveness [35]. High levels of m6A methyltransferase complex subunits suggest their potential as prognostic markers and therapeutic targets, e.g. METTL3 inhibition, may represent promising future therapy in primary tumors. Aberrant alternative splicing drives cancer development by generating diverse mRNA and protein variants, promoting tumor heterogeneity and enabling oncogenic isoforms or loss of tumor suppressor function [36].

Our IHC analysis revealed significantly higher expression of PRP19, PLRG1, and CDC5L in SACC cells, with occasional ectopic membrane localization of CDC5L. The expression of PLRG1, a key subunit of the NTC complex, correlated with BRM expression in SACC samples. We have shown that BAF155, a key subunit of the SWI/SNF CRC, directly interacts with CDC5L, a component of the NTC splicing machinery, as well as with WTAP and METTL3, members of the mRNA methyltransferase complex supporting the simultaneous occurrence of chromatin remodeling, RNA splicing, and m6A mRNA methylation in the cell nucleus.

We detected altered usage of transcripts of genes encoding NTC and m6A methyltransferase complexes. Our study demonstrated no aberrations in the levels of METTL14, but significant decreasing of WTAP and METTL3 between tumor and normal tissues. Nuclear morphometric analysis indicated that WTAP- and METTL3-positive nuclei were significantly smaller than unstained ones, and overall m6A levels were markedly lower in SACC cells comparing to healthy controls. The depletion of BRM ATPase of SWI/SNF complex in SACC cell line resulted in alteration of expression of NTC and m6A complex along with some EMT markers. This observation together with the direct binding by BRM of promoter/distal promoter regions of these genes and genes encoding SWI/SNF subunits demonstrated by ChIP experiment clearly indicated that the SWI/SNF complex may be a master regulator of the whole system controlling the abundance of NTC, m6A methyltransferase complexes and SWI/SNF CRCs, the latter by self-regulatory mechanism.

We observed in SACC samples a switch in splicing isoforms leading to significant consequences, such as alterations in the open reading frame (ORF), domain gain or loss, intron retention, changes in NMD (nonsense-mediated mRNA decay) sensitivity, or modifications in ORF length. Such aberrations could affect transcription start sites (TSS) or transcription termination sites (TTS). The global splicing event analysis in SACC samples comparing to healthy tissue indicated more than 5000 genes displaying aberration in alternative splicing. One of genes exhibiting alternative splicing variation in SACC cells is *METTL3* where we observed an overexpression and preferential usage of the main transcript. However, a second *METTL3* transcript, which is also overexpressed in SACC samples, undergoes NMD, suggesting a potential regulatory mechanism influencing mRNA stability in these cells. We detected similar alterations in alternative splicing in genes encoding key subunits of the SWI/SNF complex, including *SMARCA2*, *SMARCA4*, *SMARCB1*, and *SMARCC1*, and *PLRG1* and *PRP19* NTC subunits. In some cases, a strong expression of non-coding transcripts was observed, along with transcripts encoding distinct protein variants. These findings imply that splicing alterations may play a central role in SACC pathogenesis by reshaping protein diversity, potentially offering new targets for therapeutic intervention.

Thus, our findings suggest that impairment of the SWI/SNF complex may be a primary reason of the changes observed in SACC and they can be a result of the coordinated “snow ball” like effect accelerating multi-level deregulation of gene expression including wrong selection of alternative transcript variants (e.g. exon usage and transcript variant usage), production of pathological form of proteins or improper protein variants, neoantigens etc. all promoting development and progression of SACC.

In the most of tumors, a reduction in the amount of BRM ATPase is observed. However, in SACC, an increased expression of the BRM protein was noted [21]. Nevertheless, it is conceivable that changes in the quantity of the catalytic subunit of the SWI/SNF CRC in cancer cells primarily affect the formation and expression of neoantigens [37,38]. Considering this, blocking the BRM protein potentially could be associated with inhibiting the neoplastic process. In SACC a relatively low tumor mutation burden (TMB) was found. Whole-exome sequencing studies have reported an average of 0.31 non-silent mutations per megabase in SACC tumors, indicating a low exonic somatic mutation rate [39]. This low tumor mutational burden observed in SACC suggests limited responsiveness to immunotherapies, which are typically more effective in highly mutated tumors. Therefore, treatment strategies often focus on targeted therapies directed at characteristic genetic alterations, such as MYB or MYBL1 gene fusions [40].

Importantly, neoantigens may arise not only from gene mutations but also from aberrant RNA splicing events, including intron retention, exon skipping, or alternative splice site usage. Tumors frequently display deregulated splicing, generating tumor-specific neoantigens-an especially relevant mechanism in cancers with low mutation rates [41].

Such mechanisms may be particularly relevant in SACC, where abnormalities in the splicing machinery, along with alterations in epigenetic regulators such as SWI/SNF complexes and the epitranscriptomic m6A mRNA methyltransferase complex, may contribute to neoantigen formation despite low TMB.

Performed by us spatial transcriptomics analysis of two SACC samples in the context of BRM SWI/SNF ATPase presence-one from an early-stage tumor without nerve invasion and the other from an advanced-stage tumor with nerve invasion-revealed the heterogeneous nature of SACC, far beyond the recently published study by Yun et al. [42]. Recently published spatial transcriptomic data primarily highlighted differences between SACC cases with MYB expression and these harboring MYB–NFIB gene fusions, as well as their impact on transcriptional programs [43]. Moreover, spatial transcriptomics combined with single-cell RNA sequencing (scRNA-seq) identified upregulation of interferon-responsive genes within small HLA-positive tumor regions, suggesting that interferon signaling may reverse HLA downregulation in SACC [44] however, neither study addressed the potential contribution of alternative splicing or chromatin remodeling mechanisms in this process.

In the early-stage tumor, fewer heterogeneous and more uniform clusters were observed, whereas the advanced-stage SACC exhibited a more diverse and heterogeneous cell population. Furthermore, in the early stage of the disease, *SMARCA2* expression closely corresponded with BRM protein abundance within the same regions of the samples. The expression of other SWI/SNF subunits, including *SMARCA4*, *SMARCB1*, *ARID1A*, *SMARCC2* and *SMARCC1* genes, was predominantly localized to the same areas as *SMARCA2* transcript and BRM protein. Moreover, the expression pattern of NTC subunits *PLRG1* and *CDC5L* as well as m6A methyltransferase complex subunits *METTL14* and *METTL3* localized in the same area as SWI/SNF subunits. In advanced SACC, the expression pattern of *SMARCA2* differed from BRM protein abundance, possibly due to an alternative transcript switch and the expression of non-coding or NMD-targeted *SMARCA2* transcripts. Other subunits of the SWI/SNF complex, NTC splicing machinery, and m6A methyltransferase complex exhibit expression patterns similar to *SMARCA2* transcripts, though with varying intensities. Several of these genes, including *METTL3*, *SMARCC1*, *SMARCB1*, and *SMARCA4*, undergo alternative splicing switches. The analysis of these two clinically distinct SACC cases suggests that alternative splicing alterations likely arise during tumor progression to more advanced stages. Our findings suggest that splicing alterations during tumor progression can decouple mRNA and protein levels, highlighting the role of post-transcriptional regulation and the limited reliability of transcript data alone and support emerging hypotheses indicating such phenomenon [45]. Thus, the necessity of integrating both transcriptomic and proteomic analyses to accurately assess gene expression and its functional implications in cancer is clearly called.

In conclusion, SACC exhibits notable resistance to current therapeutic approaches. The conducted analysis unequivocally suggests that disruptions in proteins responsible for mRNA methylation, subunits of the SWI/SNF complex, and NTC are implicated in the formation and progression of SACC (Fig.6). These results not only contribute to a deeper comprehension of molecular aberrations in SACC but, more importantly, introduce novel variables for prognostic considerations. Furthermore, this findings can be important in advancing further research on the potential development of future therapeutic strategies for SACC.

**Figure 6.**
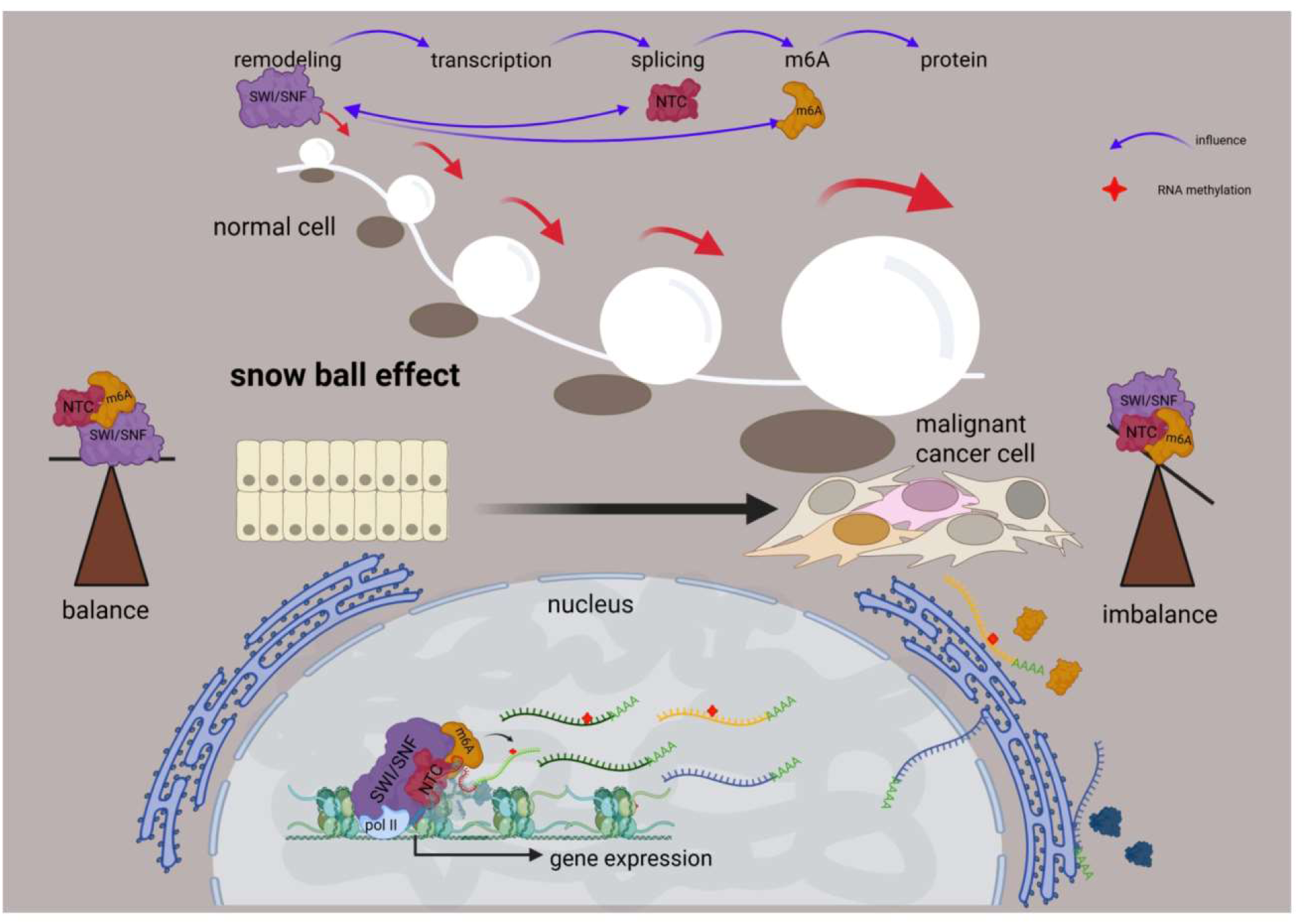
A schematic diagram illustrating the interdependence between the SWI/SNF complex, NTC splicing machinery, and the m6A methyltransferase complex in healthy cells and salivary gland adenoid cystic carcinoma cells.

## Methods

### Patient characteristic

Patients with SACC were treated in MSCNRIO. 48 patients were enrolled (29 males and 19 females) with median age 60. Primary patient samples were collected from adult patients after written informed consent in accordance with the Declaration of Helsinki. The study was approved by the Ethics Committee of the Maria Sklodowska-Curie National Research Institute of Oncology (approval No. 31/2014/1/2017). No participants received financial compensation. No exclusions were made based on race, ethnicity, sex, gender, age, or socioeconomic status. Sex was assigned according to clinical records. Owing to the exploratory nature of the study, sample size was not predetermined statistically. The details of study group are described in Table 1.

**Table 1.** Characteristics of patients of the study group. The histological assessment of SACC subtypes was performed by a pathologist.

| Variable | Value n (%) |
| --- | --- |
| Sex | n=48 |
| Female | 19 (39.6) |
| Male | 29 (60.4) |
| Age, M ±SD | 60.58 ±13.11 |
| Type |  |
| Primary | 37 (77.1) |
| Metastasis | 5 (10.4) |
| Reccurence | 6 (12.5) |
| *Histological subtype* |  |
| No type | 1 (2.1) |
| Tubular | 13 (27.1) |
| Solid | 6 (12.5) |
| Cribriform | 28 (58.3) |

### Immunohistochemistry

Paraffin-embedded tissue blocks were used, sliced and deparaffinized according to the standard protocol. Sections were incubated with particular monoclonal antibodiesː PRP19 (Abcam ab27692), CDC5L (Abcam ab31779), PLRG1 (Abcam ab86050), METTL3 (CST, E3F2A), METTL14 (CST, D8K8W), WTAP(CST, E705B) according to protocol. The EnVision® FLEX+ (Dako, GV 80011-2) kit was used for developing and horseradish peroxidase (DAB, 3,3’-diaminobenzidine) for visualization of particular target proteins in SACC samples. Hematoxylin staining was used to visualize cell nuclei. Sections were fixed by DPX Mountant for histology andvisualized using an inverted microscope. The quantification of stained cells was calculated based on the Histological Score using the following formula: H-Score = 3 × % heavily stained cells + 2 × % medium-stained cells + % poorly stained cells.

### Co-Immunoprecipitation

Co-immunoprecipitation was performed on the nuclear fraction of HeLa cells (approximately 2x10^-6^) according to the method described by [46]. Briefly, cell nuclei were lysed in ice-cold RIPA buffer (50 mM Tris-HCl, pH 7.5, 150 mM NaCl, 1% NP-40, 0.5% sodium deoxycholate, 0.1% SDS, and protease inhibitor cocktail) for 30 minutes on ice. Lysates were cleared by centrifugation at 14,000 rpm for 15 minutes at 4°C and incubated with 2 µg of specific antibodies against BAF155 (CST, 11956), CDC5L (Abcam, ab314000), METTL3 (Abcam, ab264211) or control normal IgG (CST, 2729) overnight at 4°C with gentle rotation. The antibody-protein complexes were captured using protein A/G-coated magnetic beads by incubation for 2 hours at 4°C and washed five times with ice-cold lysis buffer to remove non-specifically bound proteins. The immune-complexes were eluted by boiling the beads in 2X SDS sample buffer for 5 minutes and analyzed by Western blotting with an appropriate antibody to identify the interacting partners.

### Yeast-2-Hybrid assay

In the yeast two-hybrid (Y2H) assay, the CDS of BAF155 and CDC5L were cloned into pGBT9 and pGAD424 plasmids: one fused to the GAL4 DNA-binding domain (BAF155) and the other to the GAL4 activation domain (CDC5L). These plasmids were co-transformed either pairwise or with empty vector (control) into a Y190 yeast strain (Clontech). The direct interaction between BAF155 and CDC5L proteins reconstituted the GAL4 transcription factor demonstrated as the activation of LacZ reporter gene expression in replica lift assay performed according to the manufacturer’s protocol (Matchmaker II, Clontech).

### Microarray Analysis

The microarray datasets were retrieved from the Gene Expression Omnibus (GEO) database under accession number GSE28996 for SACC samples and GSE36820 for normal salivary gland samples. Nine SACC datasets and three corresponding files of healthy salivary gland samples were subjected to reanalysis using GeneSpring GX software (Agilent) following a standard protocol.

### BRM suppression in the SACC cell line, RT-qPCR and Western blot

The human SACC cell line UM-HACC-2A (ABM, Canada) was cultured in PriGrow III medium (ABM, Canada) with 10% heat-inactivated FBS (Biowest, France), hydrocortisone (0.4 μg/mL; ABM, Canada), EGF (20 ng/mL; ABM, Canada), insulin (5 μg/mL; ABM, Canada), and L-glutamine (2 mmol/L; ABM, Canada). Stable tetracycline-inducible shRNA targeting the ATPase (DEXDc) domain–encoding region of the SMARCA2 gene (shDEXDc) was generated using the BLOCK-iT™ Inducible H1 RNAi system (Thermo Fisher Scientific, USA). Cells were transfected with pcDNA™6/TR (Thermo Fisher Scientific, USA) via Lipofectamine™ 2000 (Thermo Fisher Scientific, USA), selected with Blasticidin (20 μg/mL; Thermo Fisher Scientific, USA), and TetR-positive clones were transfected with pENTR™/H1/TO-shDEXDc and selected with Blasticidin (20 μg/mL) and Zeocin (200 μg/mL; Thermo Fisher Scientific, USA). Construct presence was confirmed by PCR, two independent clones were identified. BRM depletion was induced by tetracycline. Total RNA was isolated using the ReliaPrep™ RNA Tissue Miniprep System (Promega, USA). 1 µg of RNA was reverse-transcribed using the Transcriptor First Strand cDNA Synthesis Kit (Roche, Switzerland) with oligo(dT) primers. Quantitative PCR was performed using iQ™ SYBR® Green Supermix (Bio-Rad, USA) according to the manufacturer’s instructions. Relative gene expression (SMARCA2, METTL3, WTAP, CDC5L, and PLRG1) was calculated using the 2(−ΔΔ)Ct method with UBC as the internal control. Data are presented as mean ± SD from three independent biological replicates. Primer sequences are provided in Supplementary Table 2. For immunoblotting, 10 µg of total protein were resolved by SDS-PAGE, transferred onto PVDF membranes (Millipore, USA) and incubated with primary antibodies against BRM (#11966, Cell Signaling Technology, USA), Vimentin (#5741, Cell Signaling Technology, USA), N-cadherin (#13116, Cell Signaling Technology, USA), WTAP (#41934, Cell Signaling Technology, USA), METTL3 (#86132, Cell Signaling Technology, USA), CDC5L (ab31779, Abcam, UK), followed by HRP-conjugated anti-rabbit IgG (#1705046, Bio-Rad, Bio-Rad, USA). Signals were detected using an Azure Biosystems imaging system (Azure Biosystems, USA). Total protein loading was assessed by 0.5% TCE staining.

### Chromatin immunoprecipitation (ChIP)

ChIP was performed according to [47], with following modifications. Chromatin was isolated from 5 × 10⁶ SACC UM-HACC-2A cells cross-linked directly in the Petri dish with 1.5 ml of PBS (pH 7.4) containing 1% formaldehyde for 4 minutes at room temperature. The chromatin was then harvested and sheared to an average fragment size of 100–500 bp using a Bioruptor® Pico sonication device (Diagenode) for 15–25 cycles (30 s on/30 s off at ultra-high power). Following sonication, the samples were treated with RNase A (125 µg/mL; Thermo Scientific) for 15 minutes on ice and centrifuged at 13,800 rpm for 10 minutes at 4 °C. The chromatin-containing supernatant was diluted to a final volume of 2 mL with wash buffer and incubated overnight at 4°C with anti-BRM (Cell Signaling Technology, #11966S) or normal control rabbit IgG (Cell Signaling Technology). Chromatin was immunoprecipitated with magnetic beads (ChIP-grade Protein G+A magnetic beads, Cell Signalling Technology; mixed at a 1:1 ratio) for 2 hours at 4°C, washed three times with wash buffer and eluted with elution buffer (10 mM Tris-HCl, 300 mM NaCl and 1% SDS). Cross-links were then reversed overnight at 65 °C. The recovered DNA was purified and analyzed by quantitative PCR (qPCR).The relative recovery of immunoprecipitated DNA was calculated using the 2^(−ΔΔ)Ct method. For normalisation, the Ct value of the mock IgG control was set to 1, and the relative fold enrichment of each sample was expressed as the fold change relative to the IgG control.

### Statistical analysis

Statistical analysis was performed using R statistical software (version 4.2.1), with the significant value p <0.05. The Shapiro-Wilk test has been used to test normal distribution. The level of the examined parameters was compared between the groups using the Kruskal-Wallis test or ANOVA when three or four groups were compared or using the Mann-Whitney U test when two groups were compared. The correlation between the variables was analyzed with Kendall’s tau-b coefficient.

### Spatial transcriptomics

All steps were performed according to manufacturer’s protocol for 10X Visium. The material for the study consisted of 2 paraffin blocks from patients with SACC. Fragments of neoplastic tissue and an area of healthy tissue were selected. In one patient, infiltration of nervous tissue occurred. Further stages of material processing were performed according to the Visium Spatial Gene Expression for FFPE CG000408 procedure using the reagents recommended by the manufacturer (10x Genomics). After verification of the material with dimensions of approximately 6.5 x 6.5 mm, the blocks were cut using a microtome into 5 µm thick slices, placed in a water bath at 43°C and transferred to a Visium Spatial Gene Expression slide (10x Genomics), dried at 40°C for 1 hour and 45 minutes. The prepared slide was deparaffinized. After this step, the slide was subjected to immunofluorescence staining using the BRM antibody ((D9E8B) Cell Signaling Technology) at a dilution of 1:200 (incubation time 1 hour RT), and the secondary antibody Alexa Fluor® 647 AffiniPure™ffiniPu₂ Fragment Goat Anti-Rabbit IgG (H+L) at 1:200 (Jackson Immuno Research, 30’). Subsequently, the slides were mounted using Ultreamount Aqueous Permanent Mounting Maedium (Daco) and scanned under a ZEISS LSM800 high-resolution microscope using software to obtain images of tissue sections and combine them to obtain a scan of the entire tissue slide. After imaging, the glycerol and coverslip were carefully removed. The slide was then dried at 37°C. The tested material was subjected to subsequent stages according to the Visium Spatial Gene Expression for FFPE CG 000407 protocol such as: Probe Hybridization Probe ligation, Probe release and extension, Visium spatial gene expression – FFPE library Construction. Sequencing and quantitative analysis of libraries were checked by the external provider (CENT, Warsaw, Poland).

### Data processing

Two samples obtained from patients with SACC were sequenced using 10x Genomics Visium technology. The data consisting of .fastq files obtained from paraffin samples along with provided microscope images was analyzed using 10x Genomics Space Ranger (v.2.1.0) The analysis utilized the reference transcriptome recommended by 10x Genomics and the Visium transcriptome probe set to generate spatial gene expression data. The processed data was then imported into the Seurat [48] (v.5.2.1) package for Single-cell sequencing analysis, where it underwent quality control (QC), normalization, and spatial clustering to identify distinct transcriptional regions within the tissue. Further analysis steps were conducted using the SPATA2 [49] (v.2.0.4) R package which was specially tailored for use with spatial transcriptomics data.

### Data denoising procedure

Employing autoencoder function available in SPATA2 original data after SCT normalization was further processed by artificial neural network. As a result of this process reduction in noise and technical artifacts was noted. The selected neural network utilized the sigmoid activation function, as it provided most consistent performance across different bottleneck neuron configurations while providing the best balance between variance retention and noise reduction. The network was trained for 20 epochs and consisted of three layers (128, 64, 32 neurons) with a dropout rate of 0.1 to prevent overfitting. All bioinformatic analyses described in this paper were conducted on both datasets. Scaled data represents the original normalized dataset and denoised data refers to the data obtained from the autoencoder.

### Clustering analysis, Differential expression, GSEA and data visualization

Clusters generated by Seurat along with manually selected healthy tissue regions, served as the basis for differential expression analysis. Most expressed genes across clusters were identified, compared and visualized using heatmaps. Expression of potential marker genes was further explored using bar plots. Gene Set Enrichment Analysis (GSEA) was performed on clusters identified by Seurat and tissue regions identified manually. Marker genes and genes of interest were also plotted across the sample to show their spatial distribution. Gene expression in both datasets was presented using violin plots. The genes of interest included those identified within the dataset as most highly expressed and associated with Gene Ontology (GO) Biological Processes (BP).

### Isoform switching analysis

Isoform switching was investigated in healthy samples of salivary gland tissue as well as SACC patients. Both datasets were obtained from the SRA NCBI database. Healthy samples were sourced from PRJNA1136833 and randomly selected (SRR29856834, SRR29856836, SRR29856840, SRR29856843, SRR29856847). Expression data from SACC patients was retrieved from PRJNA900701, including samples SRR22292847, SRR22292848, SRR22292849, SRR22292850, and SRR22292851.

Expression data was initially quantified using Salmon (v1.10.3) to prepare it for analysis with IsoformSwitchAnalyzer [50] (v2.4.0), the primary analysis tool. Data filtering was performed with gene expression cutoff of 1, isoform expression cutoff at 0 and removal of genes expressing only a single isoform. Isoform switch analysis was conducted using the recommended by the package distributors test utilizing DEXSeq (v.1.50.0). Results were annotated using GENCODE v37 annotation and .fasta files containing both protein and DNA sequences were exported.

Obtained files served as a basis for further analysis utilizing external tools. Due to the amount of sequences at this point local versions of the software had to be used. The CPC2 tool was used to determine whether detected isoforms were coding or non-coding. IUPred2 was used to identify Intrinsically Disordered Regions (IDR) and Intrinsically Disordered Binding Regions (IDBR) in proteins. Protein domain prediction was conducted using the Pfam database and the associated Perl script (pfam_script.pl). Results of these analyses were incorporated into switchAnalyzeRlist object and analysed using isoformSwitchAnalyzer package.

Alternative splicing events were analyzed to assess their biological consequences, using a dIF cutoff of 0.1. Enrichment analyses of splicing events and their consequences were performed. A volcano plot comparing relationship between dIF and isoform switch q-value was generated to highlight the presence of statistically significant and biologically meaningful switches between conditions, to further provide information about isoform switching characteristics within studied datasets another plot comparing dIF and log FC was made. Plots exploring isoforms present along with their expression patterns across the genes from SMARCA group along with METTL3 and PRPF19 were prepared. Gene ontology analysis of relevant isoforms (with consequences present) was conducted using GOTERMFINDER hosted at https://go.princeton.edu/cgi-bin/GOTermFinder.

## Data availability

Raw reads used in the study have been deposited under Bioproject accession PRJNA1426941 and are available for reviewers using the link: https://dataview.ncbi.nlm.nih.gov/object/PRJNA1426941?reviewer=2gmkg5e3kjeovl27kroehput79. Data will be published along with the article and readily available.

## Code availability

Code used for analyses described above is available at Zenodo under doi: 10.5281/zenodo.18255409 and will be made public after article release.

## Acknowledgements

The work was supported by the National Science Centre (NCN, Poland) grant No. UMO-2020/39/B/NZ5/02116 (TJS).

## Author information

K.S: Conceptualization, data curation, software, formal analysis, validation, investigation, visualization, methodology, writing–first draft, N.R: Conceptualization, data curation, software, formal analysis, supervision, validation, investigation, visualization, methodology, writing–original draft. H.S: bioinformatic analysis, visualization, methodology writing, S.K., I.P., P.C., A.B., M.J., J.K.: formal analysis, validation, investigation, visualization, methodology, M.W., S.P., A.S.,: Conceptualization, data curation, software analysis, validation, investigation, visualization, methodology, M.D., A.G., E.C., T.R., A.K., B.J.: Conceptualization, data curation, formal analysis, validation, investigation, visualization, methodology, clinical supervision of the project, P.Z: bioinformatic analysis, visualization, methodology writing, E.S., T.J.S: Conceptualization, resources, data curation, formal analysis, data and project supervision, validation, investigation, visualization, methodology, writing–original draft, project administration, writing–review and editing.

## References

1. Romani C, Lorini L, Bozzola A, Bignotti E, Tomasoni M, Ardighieri L, et al. Functional profiles of curatively treated adenoid cystic carcinoma unveil prognostic features and potentially targetable pathways. Sci Rep. 2023;13:1809.

2. Jang S, Patel PN, Kimple RJ, Mcculloch TM. Clinical Outcomes and Prognostic Factors of Adenoid Cystic Carcinoma of the Head and Neck. Anticancer Res. 2017;37:3045–52.

3. Phuchareon J, Ohta Y, Woo JM, Eisele DW, Tetsu O. Genetic Profiling Reveals Cross-Contamination and Misidentification of 6 Adenoid Cystic Carcinoma Cell Lines: ACC2, ACC3, ACCM, ACCNS, ACCS and CAC2. PLOS ONE. 2009;4:e6040.

4. Song N, Cui K, Zhang K, Yang J, Liu J, Miao Z, et al. The Role of m6A RNA Methylation in Cancer: Implication for Nature Products Anti-Cancer Research. Front Pharmacol. 2022;13:933332.

5. Cai Y, Chen K, Cheng C, Xu Y, Cheng Q, Xu G, et al. Prp19 Is an Independent Prognostic Marker and Promotes Neuroblastoma Metastasis by Regulating the Hippo-YAP Signaling Pathway. Front Oncol. 2020;10:575366.

6. Jancewicz I, Siedlecki JA, Sarnowski TJ, Sarnowska E. BRM: the core ATPase subunit of SWI/SNF chromatin-remodelling complex—a tumour suppressor or tumour-promoting factor? Epigenetics Chromatin. 2019;12:68.

7. Shukla S, Oberdoerffer S. Co-transcriptional regulation of alternative pre-mRNA splicing. Biochim Biophys Acta BBA -Gene Regul Mech. 2012;1819:673–83.

8. Cornejo KG, Venegas A, Sono MH, Door M, Gutierrez-Ruiz B, Karabedian LB, et al. Activity-assembled nBAF complex mediates rapid immediate early gene transcription by regulating RNA polymerase II productive elongation. Cell Rep. 2024;43:114877.

9. Gañez-Zapater A, Mackowiak SD, Guo Y, Tarbier M, Jordán-Pla A, Friedländer MR, et al. The SWI/SNF subunit BRG1 affects alternative splicing by changing RNA binding factor interactions with nascent RNA. Mol Genet Genomics. 2022;297:463–84.

10. Yu S, Jordán-Pla A, Gañez-Zapater A, Jain S, Rolicka A, Östlund Farrants A-K, et al. SWI/SNF interacts with cleavage and polyadenylation factors and facilitates pre-mRNA 3′ end processing. Nucleic Acids Res. 2018;46:8557–73.

11. Masliah-Planchon J, Bièche I, Guinebretière J-M, Bourdeaut F, Delattre O. SWI/SNF chromatin remodeling and human malignancies. Annu Rev Pathol. 2015;10:145–71.

12. Agirre E, Oldfield AJ, Bellora N, Segelle A, Luco RF. Splicing-associated chromatin signatures: a combinatorial and position-dependent role for histone marks in splicing definition. Nat Commun. 2021;12:682.

13. Jangi M, Sharp PA. Building Robust Transcriptomes with Master Splicing Factors. Cell. 2014;159:487–98.

14. Qiu S, Wang F, Gao X, Guan W, Dai T, Yin L, et al. Prp19/CDC5L promotes gastric cancer via activation of the MAPK pathway-mediated homologous recombination. Int J Biol Sci. 2025;21:1603–18.

15. Mu R, Wang Y-B, Wu M, Yang Y, Song W, Li T, et al. Depletion of pre-mRNA splicing factor Cdc5L inhibits mitotic progression and triggers mitotic catastrophe. Cell Death Dis. 2014;5:e1151– e1151.

16. Zhang N, Kaur R, Akhter S, Legerski RJ. Cdc5L interacts with ATR and is required for the S-phase cell-cycle checkpoint. EMBO Rep. 2009;10:1029–35.

17. Liu Z-X, Li L-M, Sun H-L, Liu S-M. Link Between m6A Modification and Cancers. Front Bioeng Biotechnol. 2018;6:89.

18. Xie H, Li J, Ying Y, Yan H, Jin K, Ma X, et al. METTL3/YTHDF2 m6A axis promotes tumorigenesis by degrading SETD7 and KLF4 mRNAs in bladder cancer. J Cell Mol Med. 2020;24:4092–104.

19. Zhao Y, Zhang L, Xia L, E H, Wang T, Lu H, et al. A METTL3-NFE2L3 axis mediates tumor stemness and progression in lung adenocarcinoma. Sci Adv. 2025;11:eadt7682.

20. Wang C, Ma D, Yu H, Zhuo Z, Ye Z. N6-methyladenosine (m6A) as a regulator of carcinogenesis and drug resistance by targeting epithelial-mesenchymal transition and cancer stem cells. Heliyon. 2023;9:e14001.

21. Jagielska B, Sarnowska E, Rusetska N, Jancewicz I, Durzynska M, Kubala S, et al. Advanced adenoid cystic carcinoma (ACC) is featured by SWI/SNF chromatin remodeling complex aberrations. J Cancer Res Clin Oncol. 2019;145:201–11.

22. Pina PSS, Jang Y, Emerick C, Scarini JF, Sousa SCOM, Squarize CH, et al. Novel Epigenetic Modifiers of Histones Presenting Potent Inhibitory Effects on Adenoid Cystic Carcinoma Stemness and Invasive Properties. Int J Mol Sci. 2024;25:1646.

23. Ferrarotto R, Mitani Y, McGrail DJ, Li K, Karpinets TV, Bell D, et al. Proteogenomic Analysis of Salivary Adenoid Cystic Carcinomas Defines Molecular Subtypes and Identifies Therapeutic Targets. Clin Cancer Res. 2021;27:852–64.

24. Tang Y-F, An P-G, Gu B-X, Yi S, Hu X, Wu W-J, et al. Transcriptomic insights into adenoid cystic carcinoma via RNA sequencing. Front Genet. 2023;14:1144945.

25. Zhu Z-M, Huo F-C, Zhang J, Shan H-J, Pei D-S. Crosstalk between m6A modification and alternative splicing during cancer progression. Clin Transl Med. 2023;13:e1460.

26. Wang S, Lv W, Li T, Zhang S, Wang H, Li X, et al. Dynamic regulation and functions of mRNA m6A modification. Cancer Cell Int. 2022;22:48.

27. Lu Y, Chan Y-T, Tan H-Y, Li S, Wang N, Feng Y. Epigenetic regulation in human cancer: the potential role of epi-drug in cancer therapy. Mol Cancer. 2020;19:79.

28. Hong J, Xu K, Lee JH. Biological roles of the RNA m6A modification and its implications in cancer. Exp Mol Med. 2022;54:1822–32.

29. An Y, Duan H. The role of m6A RNA methylation in cancer metabolism. Mol Cancer. 2022;21:14.

30. Pan J, Huang T, Deng Z, Zou C. Roles and therapeutic implications of m6A modification in cancer immunotherapy. Front Immunol. 2023;14:1132601.

31. Wang T, Kong S, Tao M, Ju S. The potential role of RNA N6-methyladenosine in Cancer progression. Mol Cancer. 2020;19:88.

32. Huang Q, Mo J, Liao Z, Chen X, Zhang B. The RNA m6A writer WTAP in diseases: structure, roles, and mechanisms. Cell Death Dis. 2022;13:852.

33. Achour C, Bhattarai DP, Groza P, Román Á-C, Aguilo F. METTL3 regulates breast cancer-associated alternative splicing switches. Oncogene. 2023;42:911–25.

34. Shi J, Chen G, Dong X, Li H, Li S, Cheng S, et al. METTL3 Promotes the Resistance of Glioma to Temozolomide via Increasing MGMT and ANPG in a m6A Dependent Manner. Front Oncol. 2021;11:702983.

35. Liu J, Eckert MA, Harada BT, Liu S-M, Lu Z, Yu K, et al. m6A mRNA methylation regulates AKT activity to promote the proliferation and tumorigenicity of endometrial cancer. Nat Cell Biol. 2018;20:1074–83.

36. Lv X, Sun X, Gao Y, Song X, Hu X, Gong L, et al. Targeting RNA splicing modulation: new perspectives for anticancer strategy? J Exp Clin Cancer Res CR. 2025;44:32.

37. Kirk NA, Kim K-B, Park K-S. Effect of chromatin modifiers on the plasticity and immunogenicity of small-cell lung cancer. Exp Mol Med. 2022;54:2118–27.

38. Tien F-M, Lu H-H, Lin S-Y, Tsai H-C. Epigenetic remodeling of the immune landscape in cancer: therapeutic hurdles and opportunities. J Biomed Sci. 2023;30:3.

39. Ho AS, Kannan K, Roy DM, Morris LGT, Ganly I, Katabi N, et al. The Mutational Landscape of Adenoid Cystic Carcinoma. Nat Genet. 2013;45:791–8.

40. Miller LE, Au V, Mokhtari TE, Goss D, Faden DL, Varvares MA. A Contemporary Review of Molecular Therapeutic Targets for Adenoid Cystic Carcinoma. Cancers. 2022;14:992.

41. Xie N, Shen G, Gao W, Huang Z, Huang C, Fu L. Neoantigens: promising targets for cancer therapy. Signal Transduct Target Ther. 2023;8:9.

42. Yuan X, Dong Z, Zhang B, Li Q, Jiang W. Combining single-cell spatial transcriptomics and molecular simulation to develop in vivo probes targeting the perineural invasion region of adenoid cystic carcinoma. Heliyon. 2024;10:e34628.

43. Ebinumoliseh I, Bijukumar G, Hoff K, Brayer KJ, Bearer EL, Ness SA, et al. Gene-Expression Programs in Salivary Gland Adenoid Cystic Carcinoma Analyzed Using Single-Cell and Spatial Transcriptomics. bioRxiv. 2025;2025.09.01.673548.

44. Li A, Gonda BL, Codd EM, von Paternos A, Mitchell DR, Herrmann MD, et al. Reversible downregulation of HLA class I in adenoid cystic carcinoma. J Immunother Cancer. 2025;13:e011380.

45. Ghoshdastider U, Sendoel A. Exploring the pan-cancer landscape of posttranscriptional regulation. Cell Rep. 2023;42:113172.

46. Sarnowska E, Kubala S, Cwiek P, Sacharowski S, Oksinska P, Steciuk J, et al. A non-canonical function of Arabidopsis ERECTA proteins and a role of the SWI3B subunit of the SWI/SNF chromatin remodeling complex in gibberellin signaling. Plant J Cell Mol Biol. 2023;115:788–802.

47. Komata M, Katou Y, Tanaka H, Nakato R, Shirahige K, Bando M. Chromatin immunoprecipitation protocol for mammalian cells. Methods Mol Biol. Clifton, N.J.; 2014;1164:33–8.

48. Hao Y, Stuart T, Kowalski MH, Choudhary S, Hoffman P, Hartman A, et al. Dictionary learning for integrative, multimodal and scalable single-cell analysis. Nat Biotechnol. 2024;42:293–304.

49. Kueckelhaus J, Frerich S, Kada-Benotmane J, Koupourtidou C, Ninkovic J, Dichgans M, et al. Inferring histology-associated gene expression gradients in spatial transcriptomic studies. Nat Commun. 2024;15:7280.

50. Vitting-Seerup K, Sandelin A. IsoformSwitchAnalyzeR: analysis of changes in genome-wide patterns of alternative splicing and its functional consequences. Bioinformatics. 2019;35:4469–71.

