## supplementary figures for "Coordinated alterations in SWI/SNF chromatin remodeling complex, mRNA splicing and mRNA methylation as hallmarks of salivary gland adenoid cystic carcinoma (SACC)"

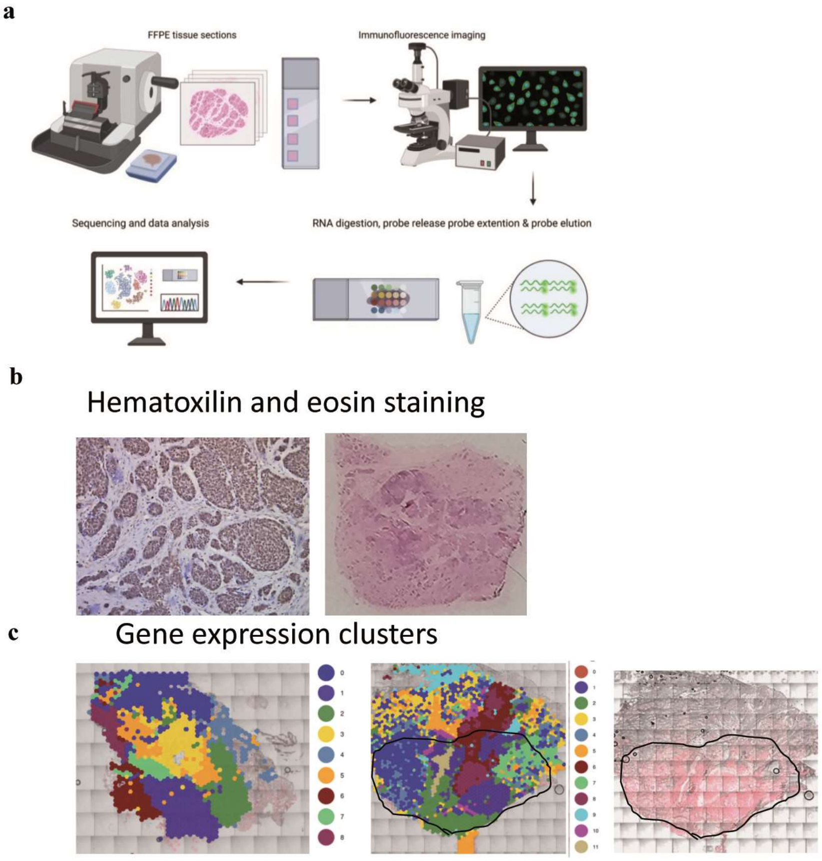


S. Fig.1 The spatial transcriptomics pipeline for SACC patients' samples analysis (a) The spatial transcriptomics schematic diagram, (b) hematoxilin and eosin staining od two SACC patient samples with and without nerve invasion (c), differentially expressed gene clusters in SACC samples.

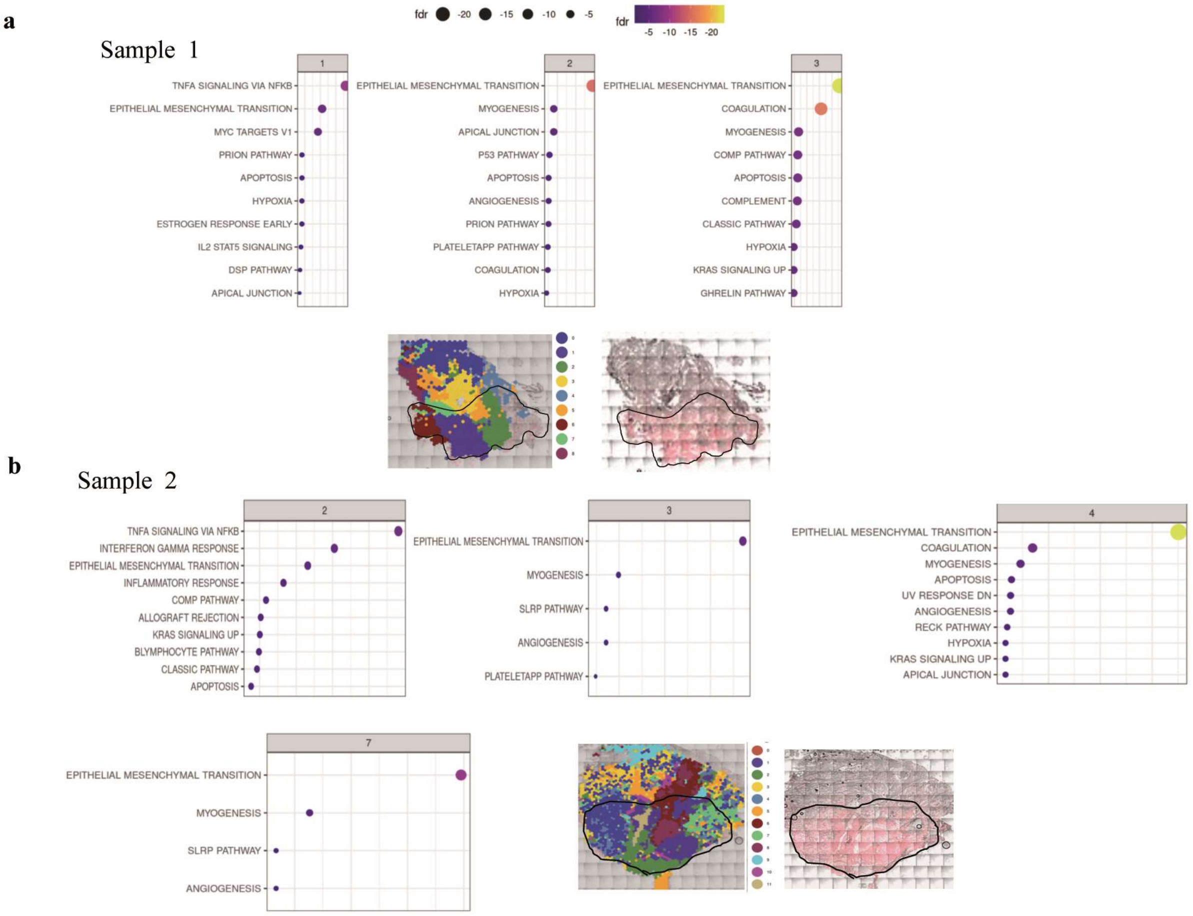


S. Fig.2 The cluster enrichment with EMT processes in sample 1 (SACC without nerve invasion) (a) and in sample 2 (advanced SACC with nerve invasion) (b) together with spatial cluster distribution and BRM protein abundance (indicated in red).

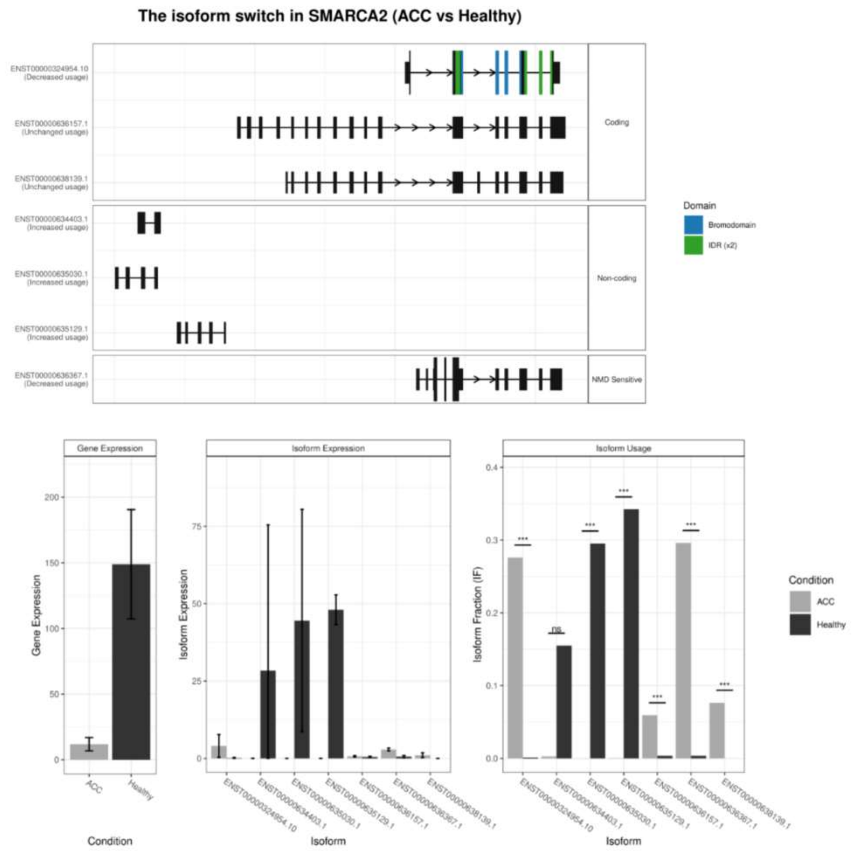


S. Fig.3 The alternative splicing aberration in *SMARCA2* gene, which encodes the ATPase BRM-the SWI/SNF complex subunit.

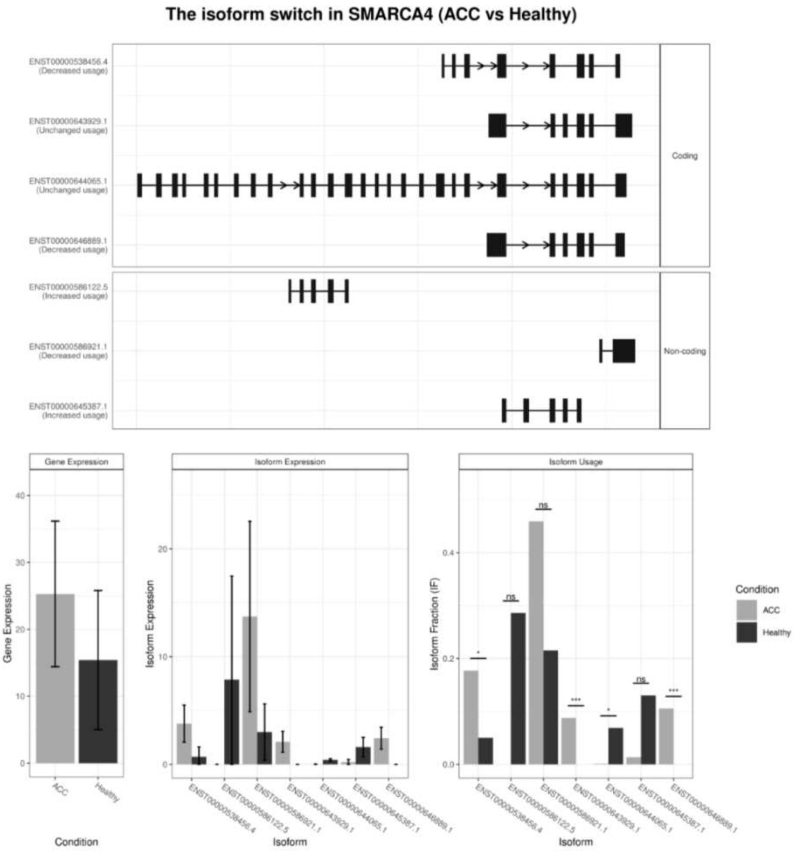


S. Fig.4 The alternative splicing aberration in *SMARCA4* gene, which encodes the ATPase BRG1- the SWI/SNF complex subunit.

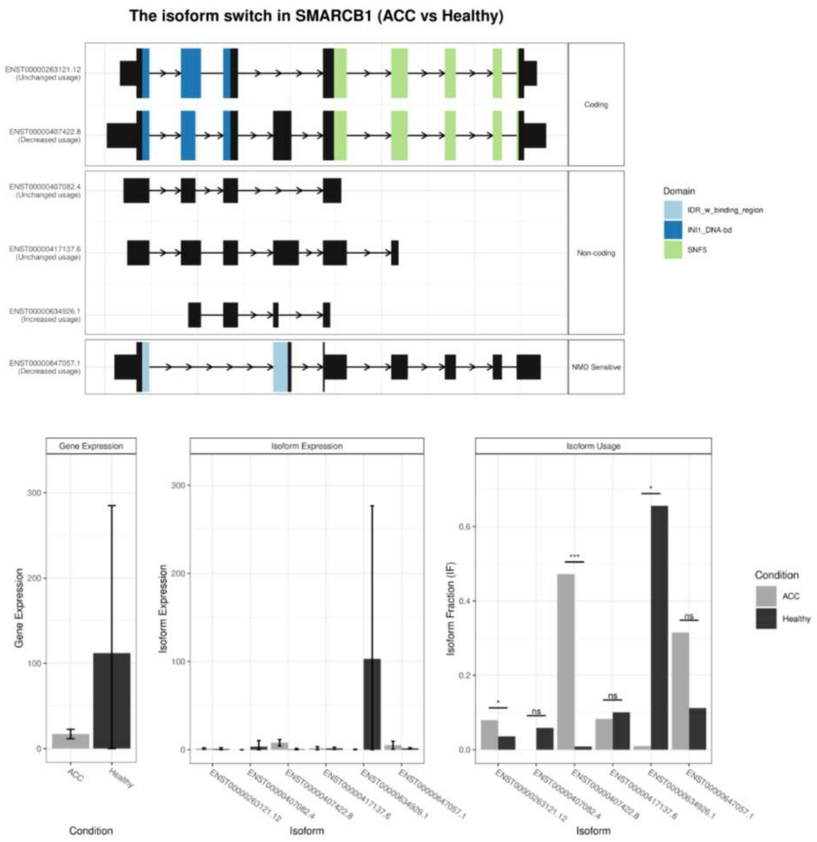


S. Fig.5 The alternative splicing aberration in *SMARCB1* gene, which encodes the one of the main SWI/SNF complex subunit – INI1.

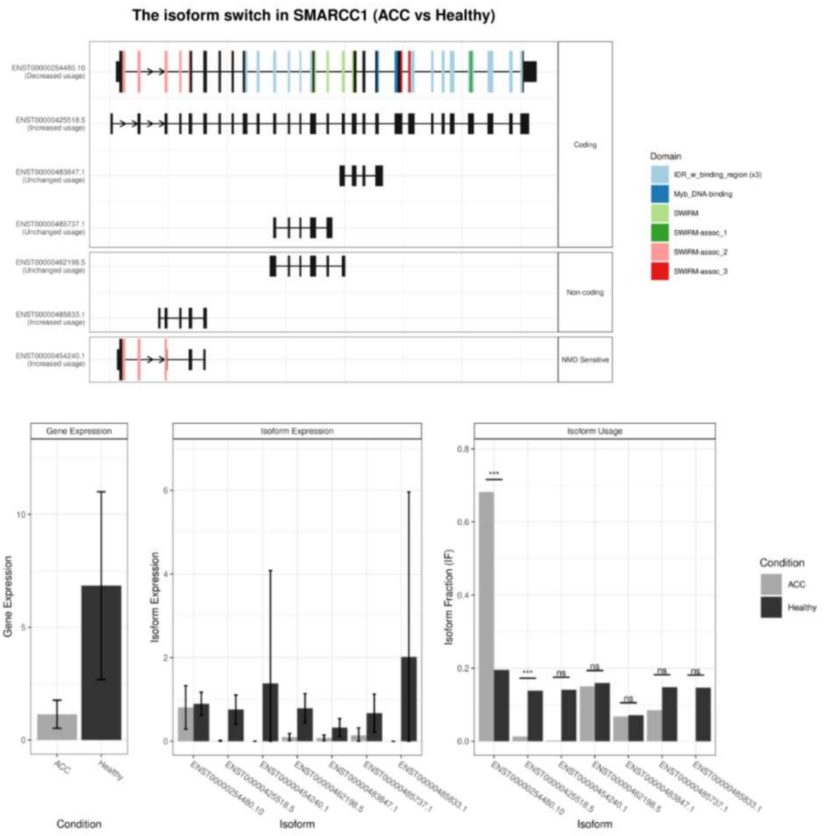


S. Fig.6 The alternative splicing aberration in *SMARCC1* gene, which encodes the one of the main SWI/SNF complex subunit – BAF155.

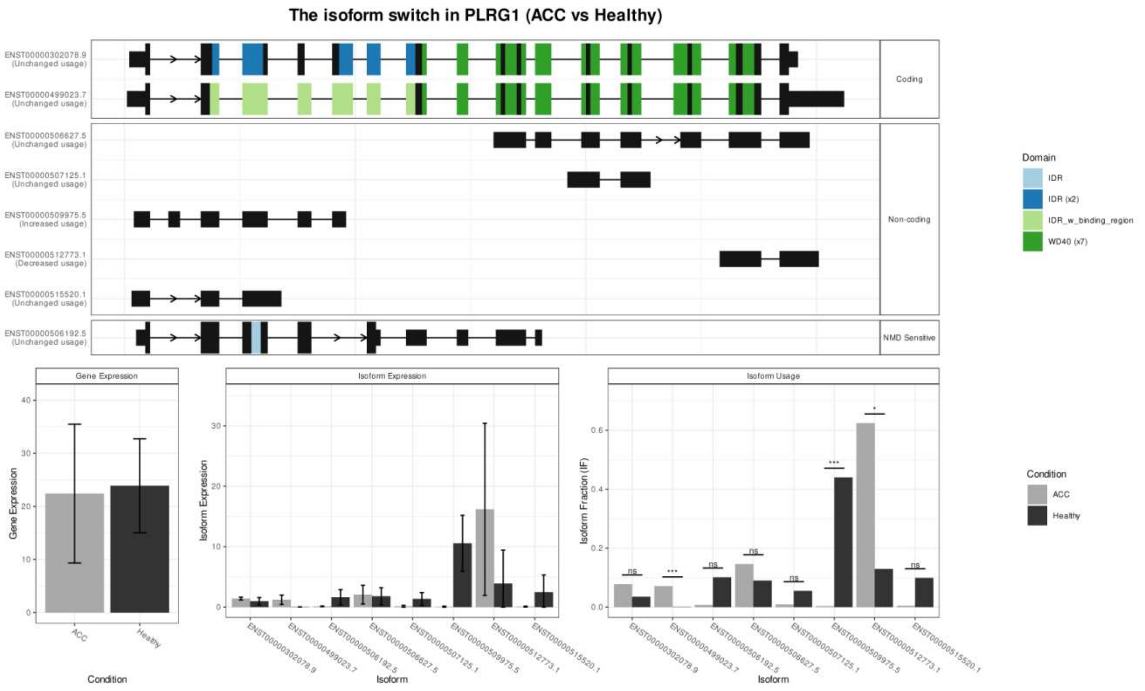


S. Fig.7 The alternative splicing aberration in *PLRG1* gene, which encodes NTC subunit.

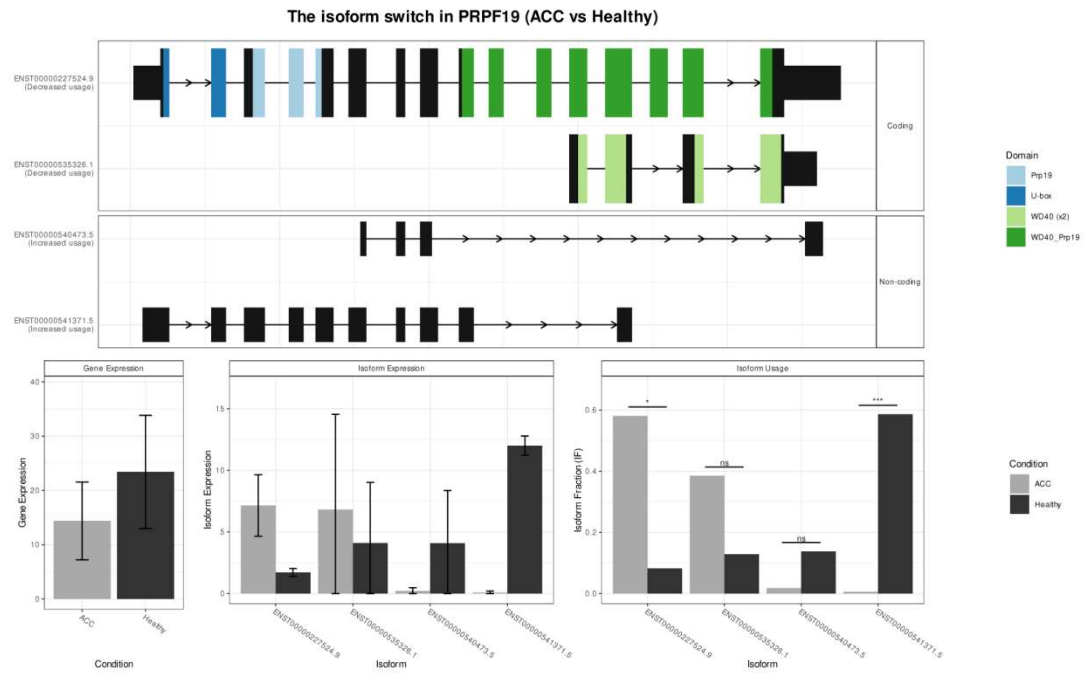


S. Fig.8 The alternative splicing aberration in *PRPF19* gene, which encodes NTC subunit.

**A**

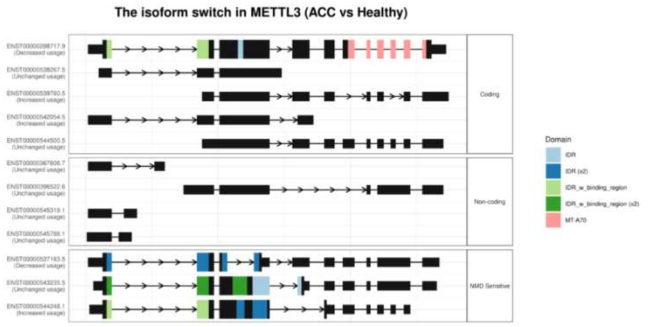


**B**

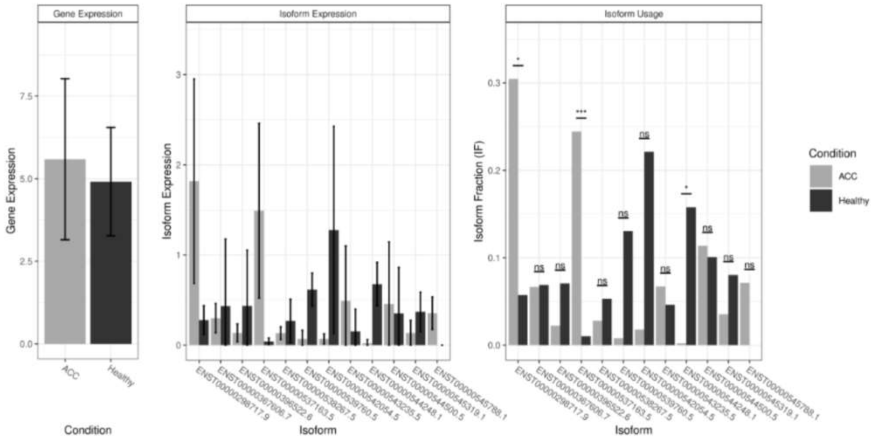


S. Fig 9 Aberrations in splicing in *METTL3* gene. (a) The METTL3 transcript isoform switch in SACC samples compare to healthy salivary gland tissue. (b) Expression pattern of particular METTL3 isoforms and their usage in SACC cells comparing to healthy cells.
